# Structural Analysis of a Metamorphic Protein using AlphaFold2

**DOI:** 10.1101/2024.11.04.621945

**Authors:** Soren Tyree, Yongick Kim

## Abstract

Metamorphic proteins, which can adopt multiple stable conformations, challenge the traditional understanding of protein structure and function. KaiB is a metamorphic protein that regulates the circadian clock, a central regulator governing gene expression in most light-perceiving organisms on Earth. An interesting aspect is that the circadian clock can be reconstituted in vitro by mixing Kai proteins (KaiA, KaiB, and KaiC) with ATP and Mg^2^+. The phosphorylation state of KaiC oscillates with a 24-hour period. The fold-switched form of KaiB binds to KaiC to activate the dephosphorylation of KaiC, while the other fold of KaiB dissociates from KaiC, allowing phosphorylation to be activated by the binding of KaiA to KaiC. To understand the metamorphic process of KaiB, we utilized AlphaFold2, a protein structure prediction program, and sequence alignments. We found that a proline residue determines the fold of KaiB. We also confirmed that mutating this proline to lysine changes KaiB to a fold-switched conformation. This validates that AlphaFold2 can be used for the study of metamorphic proteins.

## Introduction

Metamorphic proteins are unique because they can exist in two or more distinct native conformations under physiological conditions. They exhibit structural flexibility, which allows them to adapt depending on environmental factors. This ability to shift from one conformation enables them to perform multiple functions. A key characteristic is their equilibrium between different conformational states. They have multiple folds; however, they are not from protein misfolding but are functional folds that are determined by slight variations in environmental conditions. For example, a metamorphic protein may switch from a monomer to a multimer depending on its environment. This switch could be cause by a number of different factors, which sometimes can be seemingly insignificant.

There is a great interest in metamorphic proteins, especially in their evolutionary significance. The ability to adopt multiple conformations as well as perform different functions in these conformations is potentially an evolutionary advantage. This capability may be a contributing factor to the optimization of biological systems. The study of metamorphic proteins has opened up a wide range of new avenues, however, the inherent flexibility of metamorphic proteins makes the structure of all of their conformations difficult to solve. Often only one structure will be recorded in the Protein Data Bank (PDB). Therefore, predicting the alternative conformation of the folding is necessary to study the mechanism of metamorphism.

Developed by DeepMind, AlphaFold2 represents a breakthrough for protein folding predictions. The prediction of protein folding based solely on the amino acid sequence has been a sought-after challenge for decades. AlphaFold has been shown to be able to predict novel interface types which could have potential in extending definitions for existing interface types (Lee et al. 2024). Protein functions are highly dependent on their three-dimensional structure. Proteins also play a critical role in all living organisms. The advancement made through AlphaFold predictions addresses the long-standing problem and opens avenues for elucidation of many aspects, including those related to protein misfolding.

The training approach and architecture is largely the cause for the success of AlphaFold2. Many previous programs have heavily integrated our understanding of molecular driving forces into either thermodynamic or kinetic simulation of protein physics or statistical approximations, AlphaFold2 is evolutionary in which the protein structures are derived from bioinformatics analysis of evolutionary history of proteins, homology to solved structures (2021 C). AlphaFold2 takes the solved structure of proteins and their sequences that have been inputted into its data bank and compares the sequence in question and predicts the outcome based on what the folding of the solved proteins is for similar sequences across different species. This means that with the steady growth of the Protein Data Bank (PDB) the accuracy of AlphaFold2 predictions will likely increase.

AlphaFold2 introduces multiple sequence alignment (MSA) as a stage in its predictions, which septs it apart from previous programs. MSAs identify homologous proteins, which can give a better understanding for how protein structures are shaped by evolution. AlphaFold2 exploits the information in MSAs for its end-to-end training (Mirdita et al. 2022). This technique allows AlphaFold2 to generate more accurate models for the predictions for proteins with little structural data. This is especially useful for proteins which have novel folds.

AlphaFold2 has been enabled to sample alternate states of known metamorphic proteins with high confidence, however, single-sequence predictions appear to be biased toward their dominant states (Porter et al. 2023). Some alternative ways have been suggested to increase the predictive capabilities of AlphaFold2 in regards to metamorphic proteins, including the use of AlphaFold-Cluster. Although, a study this year showed that using ColabFold, an implementation of AlphaFold2, had predicted structures for proteins in three families that had higher accuracies and confidences than AlphaFold-Cluster (Porter 2024). With this study in mind, it seems that not using the MSA clusters would be a better way to predict metamorphic proteins with higher accuracy. That may not be the case. AlphaFold-Cluster predicted multiple sequences of a metamorphic protein in both conformational states and the prediction corresponded to the structure expected to be thermodynamically favored (Wayment-Steele et al. 2023).

One such metamorphic protein is KaiB, a protein in the circadian clock of cyanobacteria. The circadian clock plays a crucial role in time regulation and health maintenance in many organisms, as it is responsible for driving the circadian rhythm. The circadian rhythm allows organisms to estimate the time of day and therefore helps organisms anticipate environmental changes instead of reacting to them after they occur (Monti et al. 2018). Actually, most organisms, ranging from cyanobacteria to humans, have some form of a circadian clock (Bednarova et al. 2013). There are several reasons that understanding the circadian clock is important. Solving the underlying mechanisms could lead to medical application such as anti-aging therapies and solutions for night shift workers and others whose circadian rhythms are disrupted by their schedules. Long-term disruptions to circadian rhythms are associated with obesity, mood disorders, blood pressure issues, and an increased risk of cancer. Understanding the circadian clock’s mechanism may make it possible to regulate it and reduce or eliminate these side effects. In older individuals, the circadian clock deteriorates, leading to disruptions in sleep-wake cycles and broader physiological processes due to deficits in the amplitude and timing of the core molecular clock genes (Bednarova et al. 2018). This is what may make the circadian clock the key to anti-aging technologies. Additionally, the circadian rhythm controls the timing and extent of cell division (Martins et al. 2018). Which is why studying the circadian clock in model organisms like cyanobacteria could lead to valuable insights into reducing disruptions and shed light on certain disease processes.

Cyanobacteria was selected as the model organism for studying the circadian clock mechanism due to several advantages. Cyanobacteria was the first verified circadian oscillator that does not rely on transcriptional or translational-level regulation (Chen et al. 2017). It functions as a self-oscillating circadian clock, which is a common mechanism found in other organisms. Furthermore, cyanobacteria’s circadian clock is very simplistic, requiring only three proteins for its mechanism (KaiABC) and it can be recreated *in vitro*.

There are some requirements for the circadian clock in cyanobacteria to function, including magnesium (Mg^2+^) and adenosine triphosphate (ATP), which are need for the oscillations to occur. Mg^2+^ helps to regulate the phosphorylation and dephosphorylation of KaiC by dissociating and associating with catalytic glutamate (Glu) residues, which activate these processes (Jeong et al. 2019). Phosphorylation is linked to Mg^2+^ dissociation, while dephosphorylation is associated with its re-association. ATP is also essential to prevent KaiC from precipitating, and the ATP/ADP ratio directly impacts KaiC phosphorylation (Kim et al. 2012). Both Mg^2+^ and ATP must be added for the in vitro oscillation of the KaiABC mechanism.

Cyanobacteria’s circadian clock consists of three proteins: KaiA, KaiB, and KaiC. Recent studies have enhanced our understanding of the interactions between KaiA and KaiC, including investigations using KaiA mutations (Chen et al. 2018). KaiC, the central component of the clock, is phosphorylated and dephosphorylated by KaiA and KaiB, respectively. KaiA phosphorylates KaiC by binding to its exposed A-loop, while KaiB binds to KaiC during its phosphorylated state, causing it to dephosphorylate. KaiB is a metamorphic protein that can switch between an inactive ground state (gsKaiB) and an active fold switch form (fsKaiB) under native conditions (Tseng et al. 2017). Once KaiC is dephosphorylated, the A-loop retracts, and KaiA detaches. These three proteins, along with Mg^2+^ and ATP, can be used to reconstruct the circadian clock in vitro (Chen et al. 2018). It is essential to recreate the wild-type circadian clock oscillation in vitro to validate the lab’s techniques before introducing mutations to show credibility in the techniques used. Although research is ongoing, several mechanisms have been proposed to explain the interactions between these three proteins.

## Materials and Methods

### Structure Predictions using AlphFold2

AlphaFold2 predictions were done using 28 species of KaiB. The KaiB protein sequences were randomly collected from GeneBank at National Center for Biotechnology Information (NCBI). The protein sequences of these species were input into AlphaFold2, with predictions generated using default settings, except for two modified parameters. template_mode parameter was set to PDB 100, allowing AlphaFold2 to retrieve sequence homology templates from the Protein Data Bank (PDB), and max_msa parameter was adjusted to 32:64, following published recommendations for predicting the structures of metamorphic proteins. With each protein sequence there were five ranked models predicted, these models were either in the fold-switch conformation or the ground-state conformation. Scores were given to each of the ranked models.

If the model was a fold-switch the score given would be positive and if the model was a ground-state form the score would be negative. Rank 1 models were given a five and Rank 5 models were given a 1, and after each model of the five were accounted for a species their values were added to create the score for the species. A completely fold-switch predicted species would receive a score of +15, while a completely ground-state species would receive a score of -15.

### Investigation of the Sequence deviation using Clustal Omega

A closer inspection of the highest scored species (+15, +13) and lowest scored species (−3, +1) was done by aligning their sequences with *Thermococcus elongatus* KaiB (teKaiB) and *Synechococcus elongatus* KaiB (seKaiB) for reference. The full protein sequence for each group with identical AlphaFold2 prediction scores was entered separately into the Clustal Omega website (https://www.ebi.ac.uk/jdispatcher/msa/clustalo). Sequence similarity and deviations were then inspected manually to identify candidate mutation sites for functional study.

### Site directed mutagenesis using polymerase chain reaction (PCR)

To cause a mutation on seKaiB sequence PCR was performed. For the PCR the primers used had the desired mutation. To run the PCR 40 *μl* of water, 0.5-1 *μl* of template plasmid (between 50-100 ng), 1 *μl* of each primer (100 nM stock solution), 5 *μl* of PFU DNA polymerase buffer, 1 *μl* of PFU DNA polymerase, and 1 *μl* of dNTP were mixed into a PCR tube and then a thermal cycler was used to do 30 cycles of denaturization at 95°C for 30 sec then annealing at 53°C for 1 min and then extension at 72°C .for 8 min. Then an agarose gel was run using the sample to check the copied DNA with mutation and the desired band of around 6000 base pairs (bp) was observed then 1 *μl* Dpn1 (restriction endonuclease) was added, and the sample was incubated overnight at 37°C to remove the original template for PCR. Take 10 *μl* of PCR sample from the incubator and perform heat shock transformation at 42°C for 45 sec with the Sig10 *E. coli* strain. Then mix the resulting mixture with 1 mL of nutrient broth (2XYT) and let incubate while shaking for 1 hour at 37°C. Spin down the contents in a microcentrifuge tube and pour out the majority of the liquid. With the liquid remaining resuspend the cells and plate on an Luria Bertani (LB) and kanamycin agar plate. Then incubate overnight at 37°C. Transfer 100 *μl* of kanamycin into about 100 ml of lb broth in a flask (250 ml), then transfer 3-5 colonies with a sterile pipette tip into the broth and mix using the pipette. Put the flask into the incubator and shake, to maximize the surface area of the liquid exposed to air to make aerobic condition, overnight at 37°C. Then take 5 mL and spin down in the centrifuge at 4000×g for 15-20 minutes. Remove all liquid. Resuspend cells following the plasmid mini kit (Zymo Research) instructions. Take the plasmid sample and use the nanodrop to measure the concentration in ng/mL. Final DNA sequence was confirmed by Sanger sequencing by reading full insert sequence using T7 terminator sequencing primer

### Protein Expression and Purification

#### Expression of Kai proteins

To express Kai proteins in pET41a(+) vector, use 1 *μl* of pET-41-seKaiB or pET-41-seKaiC plasmid and add to *E. coli* BL21DE3 strain (BL21 is the *E. coli* cell and DE3 is the virus promoter). Since the plasmid is not very large (under 10k base pairs (bp)) heat shock transformation can be used. Heat shock by placing the microcentrifuge tube with the contents in a water bath heated to 42°C for 45 seconds then put on ice. Add contents to 1 ml of 2XYT and then add everything to a cell culture tube with a cap to incubate for thirty minutes to an hour at 37°C. Plate on a Luria Bertani (LB) and kanamycin agar plate using 50 *μl* of sample. Then incubate the overnight at 37°C (16-20 hours). This is a selective process, nothing but the desired *E. coli* should grow. Transfer 100 *μl* of kanamycin into about 100 ml of LB broth in a flask (250 ml), then transfer 3-5 colonies with a pipette into the broth and mix using the pipette. Put the flask into the incubator and shake, to maximize the surface area of the liquid exposed to air, overnight at 37°C. Add 1 ml of kanamycin to a large flask (2 L) with LB broth (1 L) and add all of growth sample from the 250 ml flask. Put the large flask in the incubator and shake (at 37°C for one hour, then take 1 ml and put into a visible-spectrum cuvette and measure the optical density (O.D.) at 600 nm, this is to get a rough estimate of the cell growth. When O.D. reaches 0.4, place back into the incubator at 25°C. Test the O.D. every thirty minutes has passed and once an ideal O.D. is reached (0.6-0.8) add 1 ml of Isopropyl β-D-1-thiogalactopyranoside (IPTG) (final concentration is 1 µM). Place in the incubator and shake, at 25°C overnight (16-20 hours). It is important that the temperature is correct, at higher temperatures *E. coli* does not fold the Kai proteins correctly.

#### Purification of Kai proteins

Start by separating the contents of the large flask into three bottles and place them into the centrifuge and spin down, then remove the supernatant. Resuspend the cell with 60 ml of buffer (150 mM NaCl, 50 mM Tris, 5 mM MgCl_2_, 1 mM EDTA, 1 mM DTT, pH=7.3) and split into two 30 ml tubes with screw caps and put on ice. In the case of KaiC, 1 mM ATP should be added in the buffer for entire purification process to prevent precipitation. Use sonication to break up the cells for 30 seconds and rest for 1.5 minute four times. Then make the two tubes evenly weighed and centrifuge for 45-60 minutes at 20000×g. Filter the supernatant and put sample into a GST column with a syringe at 1 ml per minute after the GST column has been washed with the buffer. The proteins have a GST tag from the plasmid used that bonds to the GST column. wash and elute the bound protein using the fast protein liquid chromatography (FPLC). For the elution buffer, 10 mM glutathione should be added. 100% elution buffer was used to elute out the bound Kai Protein. Test the concentrations using Bradford reagent, if it changes color to blue then proceed, if not concentrate using reverse osmosis (cutoff is10 kDa) tube in the centrifuge and spin for 5-20 minutes depending on concentration needed. Cut the GST tag from Kai protein by using a protease (RV3C) overnight. Then use a desalting column to do a buffer exchange for removing glutathione. Use the GST column again to remove the tag from the protein sample. Once the tag is removed use the desalting column again to exchange the buffer for the reaction buffer that is 150mM NaCl, 20mM tris and pH=8.0. Test the concentration using the Bradford assay at 595 nm absorbance. Protein concentration was determined using a standard curve generated with Bovine Serum Albumin (BSA).

### *In vitro* Oscillations

With the proteins that have been purified, the *in vitro* oscillations were performed. The oscillatory phosphoryl-transfer reaction was done using seKaiB or its mutants with the wild-type KaiA and KaiC with ATP, MgCl_2_. These reaction mixtures were put into an automatic sample collector that would collect a small sample every two hours starting with hour 0 for 48 hours. The samples were put into microcentrifuge tubes that contained 2 *μl* of SDS-PAGE loading dye (100 mM Tris·HCl, pH 6.8, 4% SDS, 0.2% bromophenol blue, 20% glycerol, 400 mM β-mercaptoethanol). These samples were loaded into the wells of 6.5% polyacrylamide gels. Using gel electrophoresis, the samples were run for 30 minutes at 60V and then 1 hour and 45 minutes at 140V while on ice. Once the gels were run, the gels were removed from the glass covers and placed into containers with Brilliant Blue Coomassie staining solution overnight. Then the gels were destained overnight in water so that only the proteins were stained. Images were taken of the gels afterwards with a white light background and used to perform the densitometry to calculate the phosphorylation of the samples at different times to observe the oscillations using ImageJ and excel.

## Results and Discussion

To classify the tendency for fold-switched KaiB based on sequence variation, we collected random KaiB sequences from GenBank (NCBI) and all available KaiB sequences from the Protein Data Bank (PDB). Each sequence was analyzed using the AlphaFold2 website, with parameters adjusted according to previously reported recommendations. We found that 5 out of 28 species showed a high tendency to form a fold-switched KaiB structure (Fig. 2).

**Figure 2:**
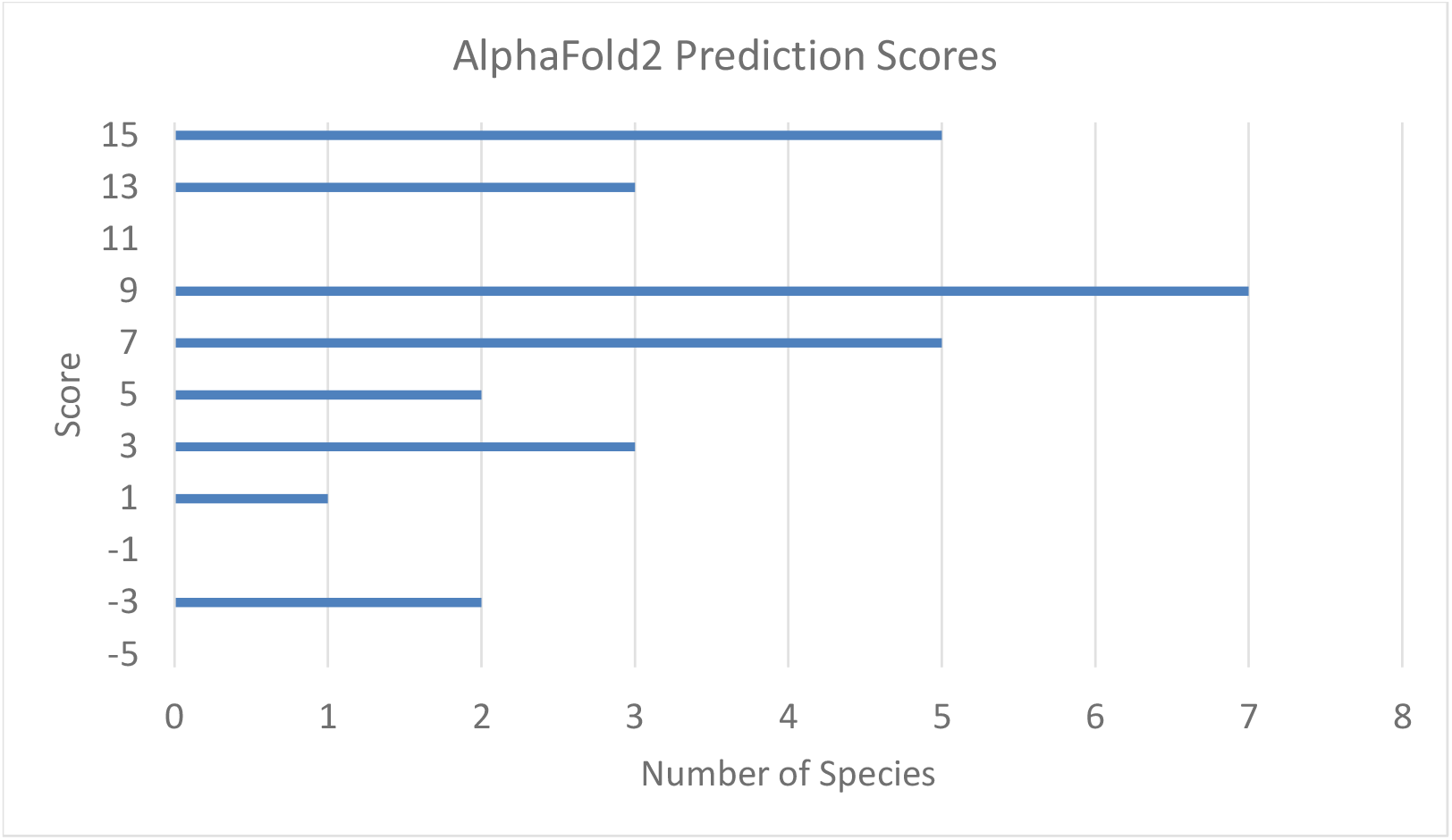
Scores and the number of species that received them from the AlphaFold2 predictions.

To analyze sequence variation among predicted fold-switched KaiB proteins, we used the Clustal Omega tool on the EMBL-EBI website. Our examination of the C-terminus of KaiB proteins revealed that most have a shorter C-terminus compared to KaiB from *Synechococcus elongatus* (seKaiB). Notably, KaiB from *Legionella pneumophila* (lpKaiB) has the shortest C-terminus, and its X-ray crystal structure in the PDB exhibits a fold-switched KaiB conformation (Fig. 2). Interaction in the C-terminal region is thought to be crucial for circadian rhythm generation and for tetramer formation, typically observed in the ground-state KaiB conformation, which does not bind to KaiC or KaiA. Additionally, previous studies reported that removing certain residues in the C-terminal region disrupts circadian rhythms in cyanobacteria (Iwase et al. 2005).

To test the effect of the C-terminus on circadian rhythms, we reconstituted *in vitro* oscillators and measured their oscillation properties. We generated a KaiB mutant with only 92 residues (seKaiB L92ter) instead of the full 102 residues (seKaiB WT). We anticipate that this shortened C-terminal mutant may have a reduced tendency to form tetramers, with more of the protein existing as monomers or dimers, which may facilitate a conformational shift toward the fold-switched form. The KaiB L92ter mutant disrupts circadian oscillation, lengthening the period by approximately four hours and reducing amplitude by around 50% compared to seKaiB WT (Fig. 4). However, it still maintains distinct oscillation patterns that were not observed in previously reported fold-switched seKaiB D90R mutants (Chang et al. 2015).

**Figure 3:**
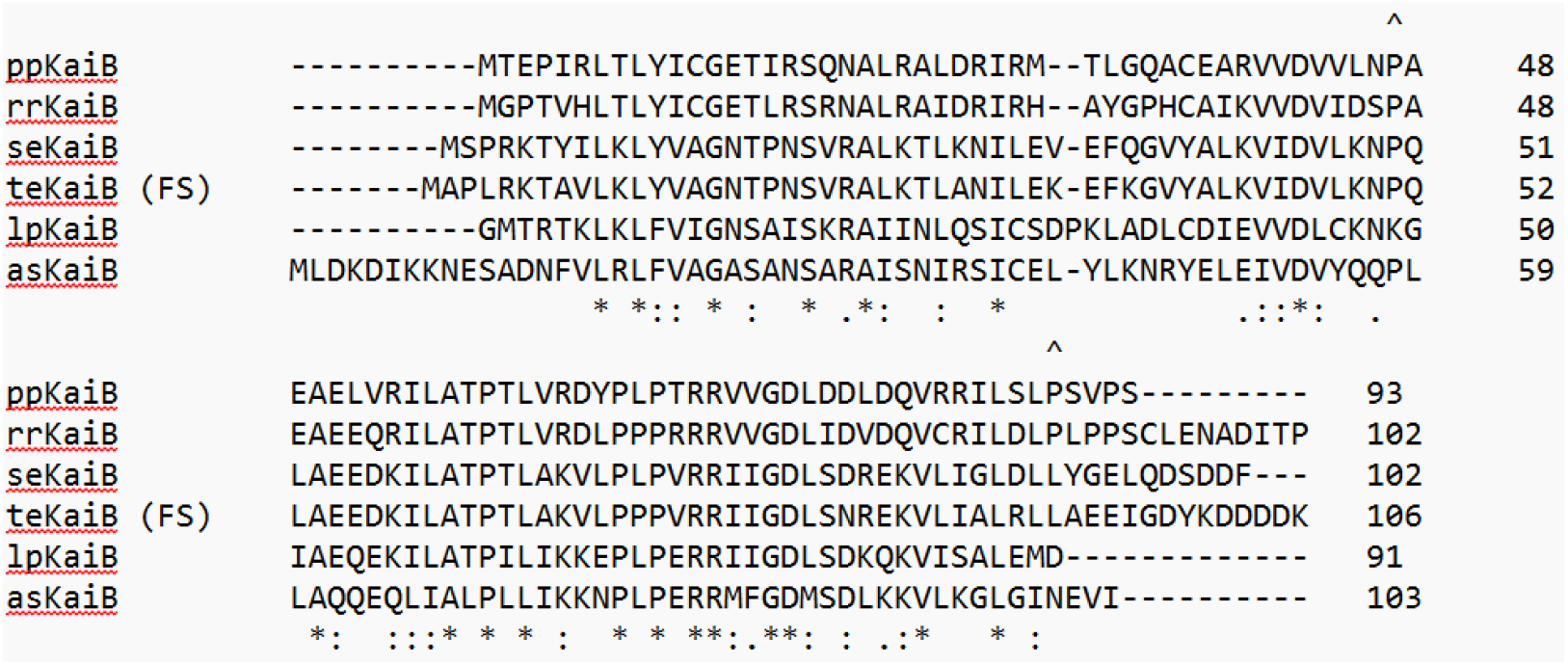
Sequence alignment of five potential fold-switched KaiB proteins with seKaiB.

**Figure 4:**
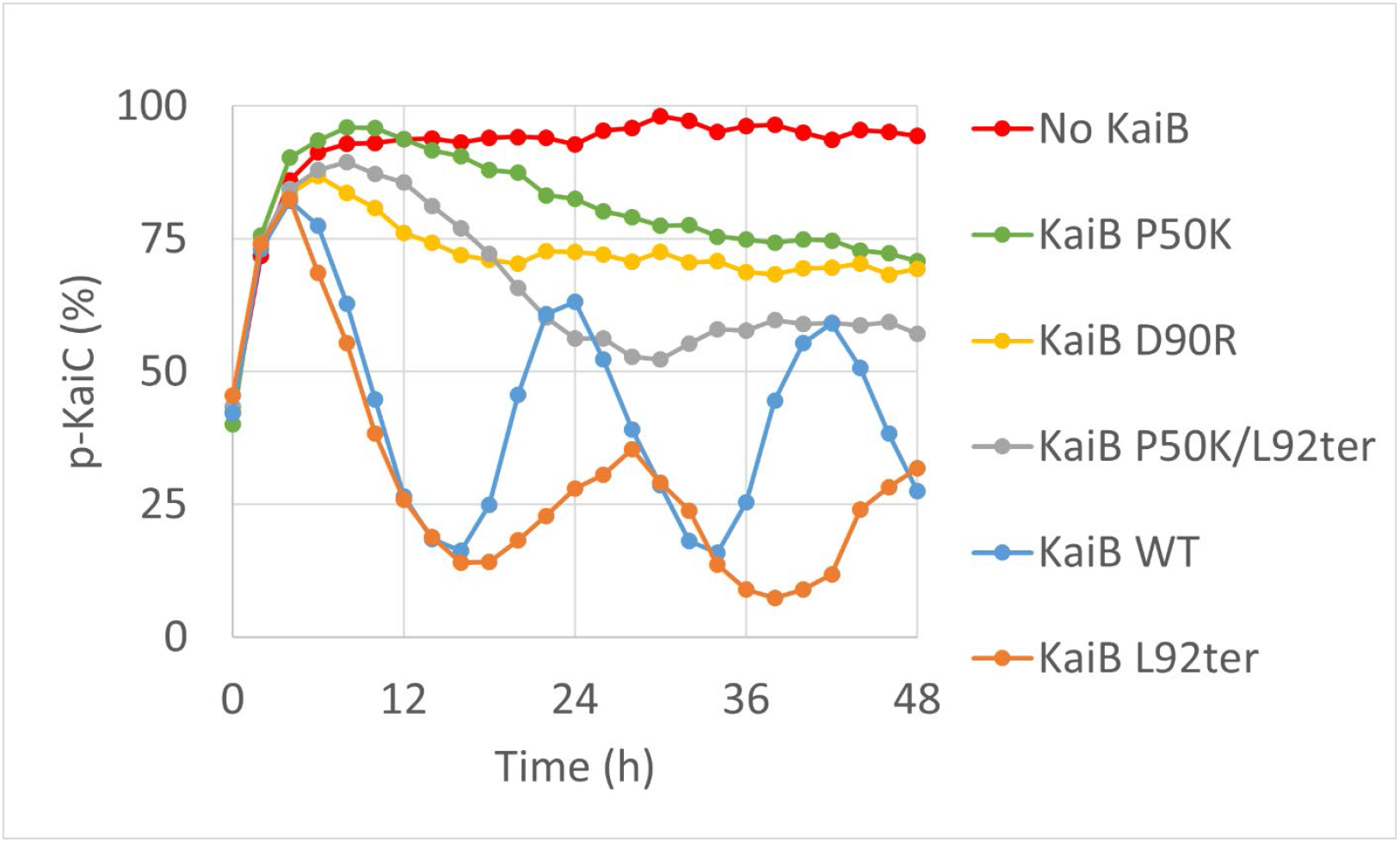
*In vitro* oscillation of KaiC phosphorylation with various seKaiB mutants.

We further investigated sequence variation and found that Pro50 (hereafter, numbering follows the seKaiB sequence) is replaced by Lys in lpKaiB (Loza-Correa et al. 2014), and Pro18 is similarly substituted with Ala (Fig. 3). We focused on proline variations because previous studies suggest that proline isomerization is key to the transition from the ground-state to fold-switched KaiB conformation (Wayment-Steele et al. 2024). To visualize these differences, we compared the ground state seKaiB structure (PDB ID: 4KSO) with the AlphaFold-predicted fold-switched KaiB structure (Fig. 5). Notably, the most significant structural deviation begins at Pro50, where an α-helix forms after Pro50 in the predicted fold-switched seKaiB, but in the ground-state structure, this region shifts to unstructured turns (Fig. 5). This suggests that helix formation and disruption may drive the interconversion between these two conformations.

**Figure 5:**
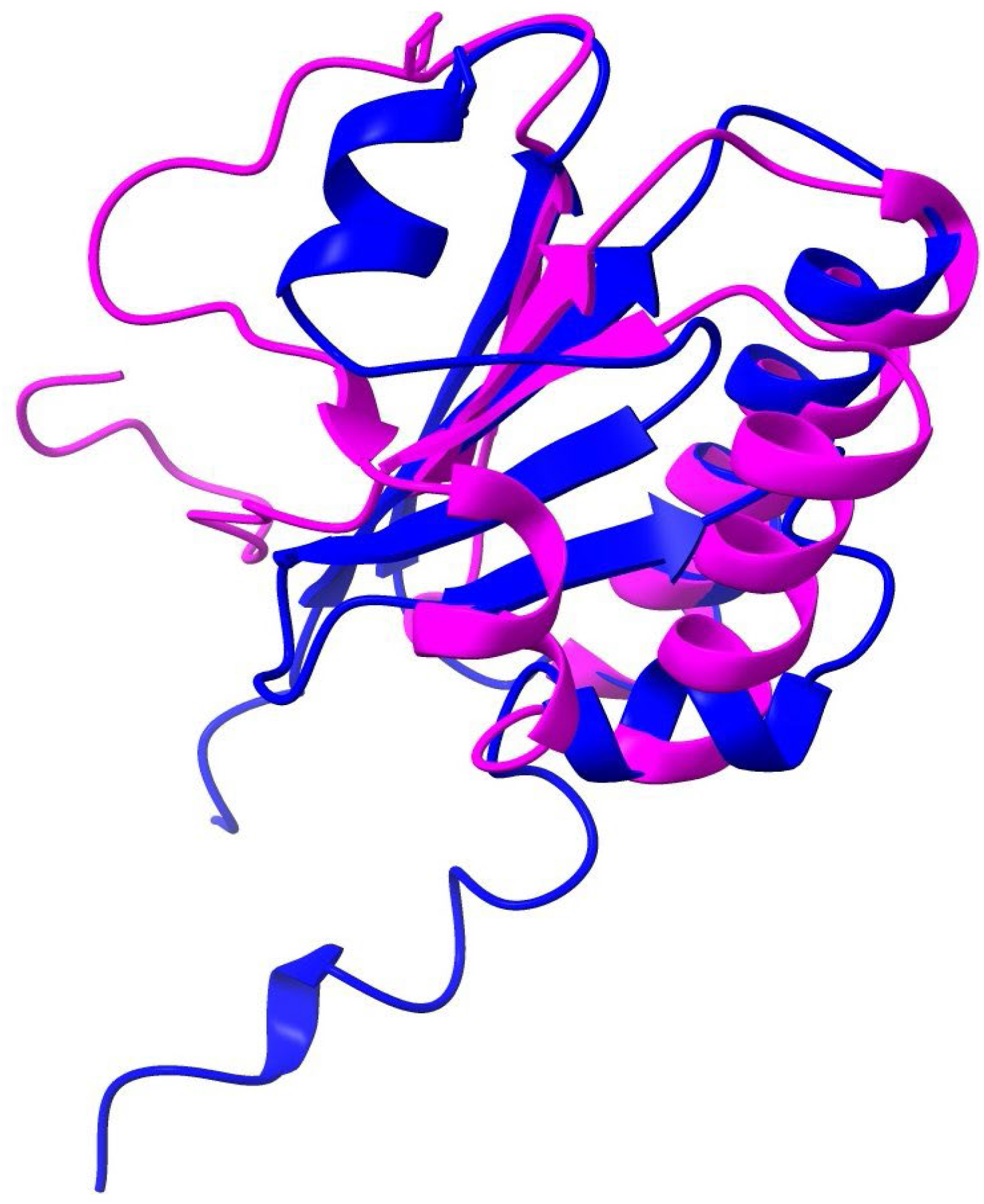
Structural comparison of ground-state and fold-switched KaiB conformations.

**Figure 6:**
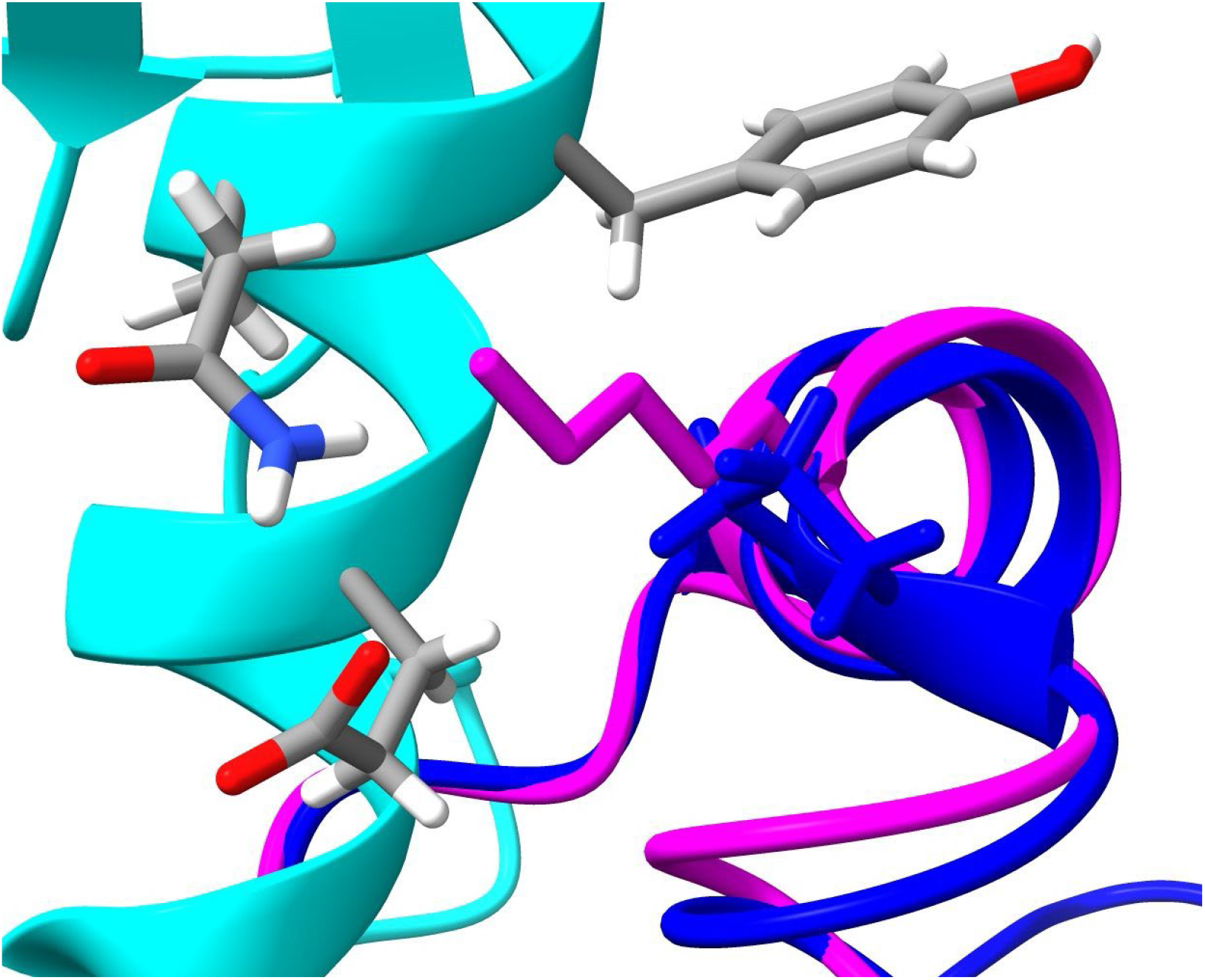
The binding interface between KaiB (Blue: seKaiB WT, Magenta: seKaiB P50K) and KaiC (Cyan).

To test this, we introduced the P50K mutation in seKaiB, replacing Pro with Lys to promote α-helix formation by removing Pro, known as a helix breaker (Li et al. 1996). We generated two P50K mutants: one with a full-length C-terminus (102 residues, seKaiB P50K) and a shortened version (92 residues, seKaiB P50K/L92ter). Both mutants lack oscillation, similar to fold-switched KaiB, yet still exhibit hyper-phosphorylation, with phosphorylation levels exceeding 50% (Fig. 4).

To understand this hyper-phosphorylation with P50K mutants, we examined the KaiB-KaiC binding interface and found that our mutation site lies within the binding region, where Lys creates steric hindrance because it has bulky side chain. This likely reduces the binding affinity between KaiB and KaiC, as observed for lpKaiB and lpKaiC previously, which also exhibit no binding affinity (Loza-Correa et al. 2014). Reduced KaiB-KaiC binding may be the primary cause of hyper-phosphorylation, as KaiB binding is crucial for initiating KaiC dephosphorylation; without this binding, KaiC remains phosphorylated.

**Table 1:**
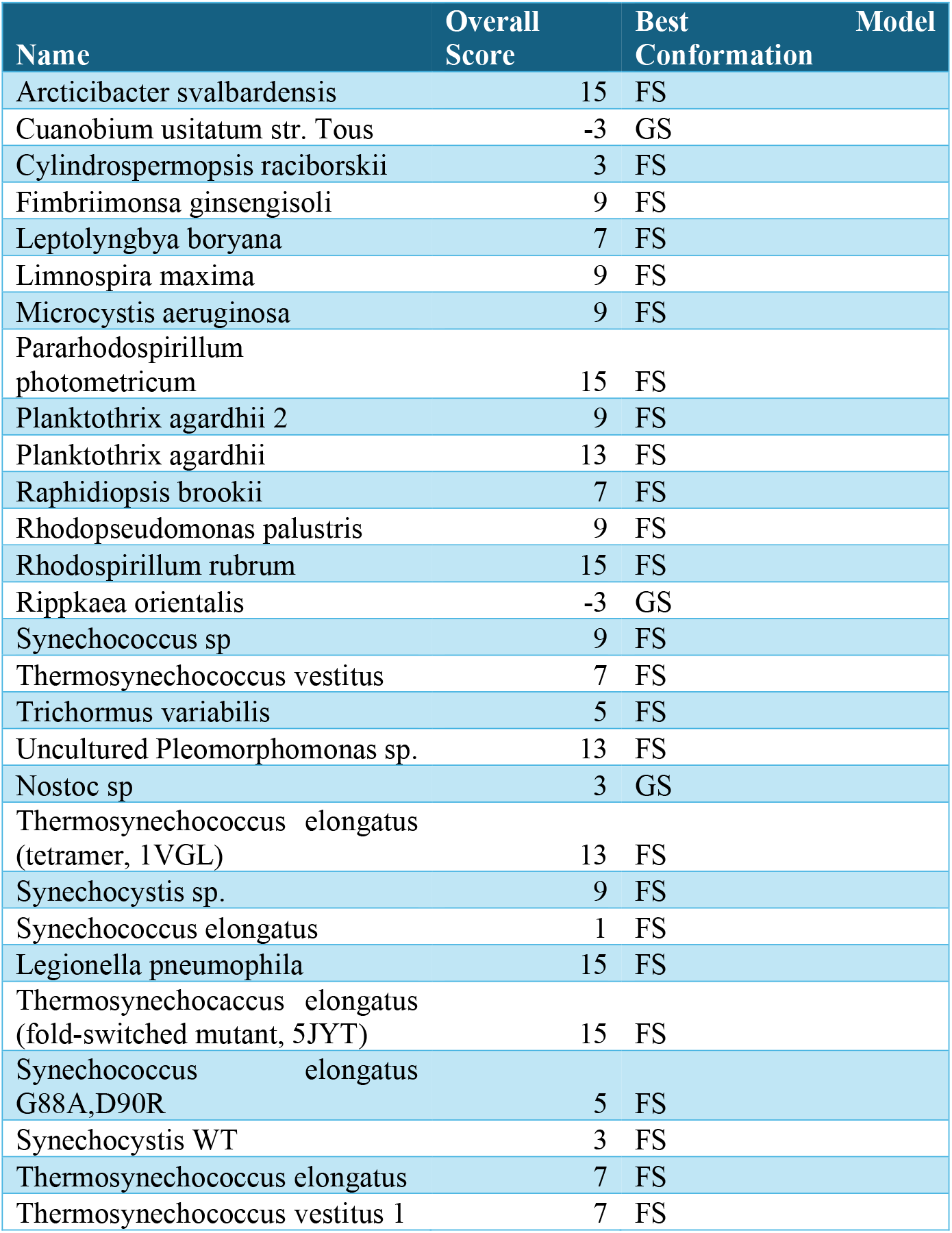
Individual Scores for each species used.

## Author Contributions

The manuscript was written through contributions of all authors. All authors have given approval to the final version of the manuscript.

## Notes

The authors declare no competing financial interest.

## ACKNOWLEDGMENT

We thank to Eugene Kim for his assistance. This research was made possible by the NASA Established Program to Stimulate Competitive Research, Grant # 80NSSC22M0027.

## Notes

### Competing Interest Statement

The authors have declared no competing interest.

